# Colorectal cancer as a model for biological evolution

**DOI:** 10.1101/2023.04.13.536757

**Authors:** Marco Ledda, Alessandro Pluchino, Marco Ragusa

**Affiliations:** Department of Physics and Astronomy, University of Catania, Catania, Italy; Department of Physics and Astronomy, University of Catania and INFN Catania division, Catania, Italy; Department of Biomedical and Biotechnological Sciences—Section of Biology and Genetics, University of Catania, Catania, Italy

**Keywords:** Agent-based Models, Colorectal cancer (CRC), Biological evolution

## Abstract

*Complexity in cancer research has led to the creation of powerful analytical tools for helping experimental* in vivo *and* in vitro *methods. These tools range from systems of differential equations resolved in computer simulations, to lattice models and agent-based models (ABMs). These analytical methods are focused on studying cell behavior and dynamic cell populations. Among these, those that are increasingly used are ABMs because they can incorporate multi-scale features ranging from the individual up to the population level, mixing rules based on mathematical and conceptual parameters, and combining statistical/population assumptions with individual heterogeneity. In this work, we present an ABM that simulates tumor progression in a colonic crypt, with the aim of providing an experimental* in silico *environment for testing results achieved in traditional lab research, and developing alternative scenarios of tumor development. As the first part of an ongoing project, the long-term goal is to reproduce and study, the general evolutionary mechanisms of a biological system*.

## 1. Introduction

The construction of a generalized model that simulates tumor development in any tissue of our organism is an undertaking that presents considerable difficulties in various aspects: there are analytical and computational aspects about availability, modelization and organization of data, and issues concerning complexity and irregularity of the genomic and histological characteristics of the various tumors [1]. These difficulties had led to prefer a particular analysis of the various tumors, which are analyzed as quasi-isolated phenomena, with their respective peculiarities regarding the function of the cells, their phenotypes, and their genotype [2]. Of all cancer types colorectal cancer (CRC) is the only one for which there is a clear developmental model, which describes the phases from normal-neoplastic epithelium to metastasis as a function of the activation and suppression of a series of oncogenes and tumor suppressor genes, respectively [3].

The theoretical model of CRC progression was developed using statistical data about genetic mutation frequencies and chromosome deletions, and further discoveries demonstrated that there are three genetic agents present in every human tumor: oncogenes, tumor suppressors and stability genes [4]. Other important genomic features of CRC progression are known as CIN (*chromosomal instability*), MSI (*microsatellite instability*), DNA mmr (*DNA mismatch repair*) and DNA methylation, which is the most studied epigenetic cause of tumor development [5]. The oncogenic multi-step process starts when there is a deregulation of the *β*-catenin pathway due to **APC** mutation [6], and proceeds with other somatic mutations that make cells more proliferative (**Kras** enhanced function) and insensible to apoptosis (**TP53** loss) [4].In this ideal timeline, the various genomic deregulations listed before occur with different frequencies [7].

The effort made to describe, classify, and predict the origins and evolution of CRC, has produced a considerable amount of experimental tools such as *in vivo* [8], *in vitro* [9, 10] and *in silico* models [11, 12]. All these approaches have their own qualities and limitations. In vivo animal models, for instance, provide strong therapeutic evidence controlling only few variables, however, even supposing the homology of the genes controlled, it is not sure if the therapy will act in the same way in human beings[13]; *in vitro* models allow observation and control at the cellular level, directly observing in real time what is happening, however, to have reliable results there must be a standardization of some cellular features such as the timing of the cell cycle; furthermore, colonic flora is absent [9, 13]; *in silico* models show a similar problem because they use compartment models and the dynamic between these compartments is regulated by ordinary differential equations that allow the analysis of continuum variables but cannot grasp cellular heterogeneity [14]. A kind of *in silico* model is called ABM (*agent-based model*), which adopts a completely different conception of how to simulate a phenomenon. The aim of an ABM is to reproduce features and the individual behavior of every agent of a particular system, and give them few rules that encode their (inter)actions, thus observing what dynamics emerge from these interactions, and if the dynamics are, in some way, coherent with real data [15, 16, 17, 18, 19, 20].

Over the last twenty years, the growing power of computers has produced a great variety of computational methods, such as lattice models and cell-based models, which can simulate the major characteristics of cancer, such as hypoxia, angiogenesis, drug-delivery, cancer stem cells, immune system interactions and cancer cells invasion of adjacent tissues and metastasis [21].

*Anoikis*, for instance, is a very well-studied phenomenon, and a good way to simulate it is with lattice models where each cell has its own space and when, for different reasons such as mutations that influence inter-cellular adhesion, the pressure between them exceeds a given threshold representing the distance from the basement membrane, the detached cells die [22, 23]. An interesting extension of the mono-crypt models [22] is the one that extends them to incorporate the proliferation in several crypts [24]. Another approach considers the comparison between cell based spatial models, incorporating cell-cell adhesion and proliferation changes in function of Wnt signals, and a stochastic one-dimensional model predicting the evolution of monoclonal conversion [25]. Another one, that focuses on genetics, considers the cell population inside the crypt, as a population evolving after changes in genotype frequencies. The dynamic of the model is studied comparing genomic analysis on real data and an agent-based model where sub-clones are generated after the acquisition of driver mutations [26].

Following the ideas developed in these models, but with a different theoretical approach, the aim of this work was to build an agent-based model, that simulates cell dynamics in the colonic crypt, to provide an experimental environment useful to test different scenarios regarding CRC development. Furthermore, we propose an approach that unifies genetic triggers, spatial and population constraints and other species interactions (immune system like cells). The model is also conceived to represent a minimal biological evolutionary system that considers cells in the crypt microenvironment as individuals belonging to different populations and species subjected to evolutionary mechanisms [27].

The rest of the paper is organized as follows. Section 2 provides a brief description of the phenomenon from a biological perspective and the model features; in Section 3 we describe and explain the simulation results; in Section 4 we discuss the implications of the results; in Section 5 we summarize the work and outline future intents.

## 2. Methods

### 2.1. Biological background

Colonic crypts are invaginations of the colon tissue where cells, called enterocytes, after being generated by stem cells located in the bottom of the crypt, move upward in the crypt to reach the crypt mouth coming out into the lumen. During such constant movement, these cells receive Wnt signals from mesenchymal cells and these signals stimulate the production of *β*-catenin, which is produced in a descendent gradient from the bottom to the top of the crypt [2, 28]. Cells with high levels of *β*-catenin are stem-like cells called *Transit-amplifying* cells, which proliferate the most, providing a constant flow of new cells in the crypt; moving upward, the level of *β*-catenin decrease allowing the cells to assume a function becoming *Differentiated* cells, entering apoptosis in 3-5 days [2, 29].

This process constitutes a remarkable mechanism for avoiding cell dysfunctions due to DNA damage or other problems caused by the intestinal microenvironment [2].

### 2.2. The model

The model is designed with the ABMs software Netlogo [30]. The interface shows the world, a 2D representation of the colonic crypt arranged as if the crypt is an unfolded cylinder.

**Figure 1.**
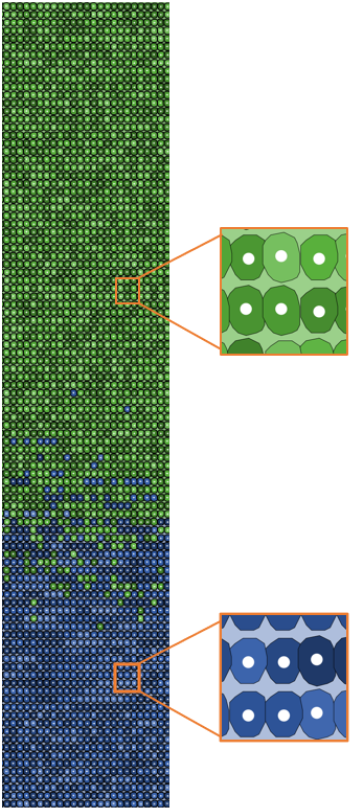
Cell typologies: blue cells *Transit-amplifying*, and green cells *Differentiated*.

### 2.3. World Structure

The size of the population we have chosen corresponds to the median crypt, which can have 2000-2500 cells with a diameter of 25 cells per line; the epithelium is renewed every 3-5 days, which means from 72 to 120 hours for every cell to either die or reach the crypt mouth [31]. The crypt height in the model has a measurement corresponding to the number of cell lines, that is 100 cells, chosen because cells make a movement forward every time step, set up to 1 hour (1 tick of the program). Considering a cell that does not die during the migration with a life-time equal to 0 in the first line and 100 in the last line, we have a maximum migration time almost corresponding to the real one [32, 33].

The crypt-world is filled with cells of the two typologies in a defined proportion that is roughly 75% *differentiated* cells, 25% *transit* cells [22]. Given that the crypt is conceived as an unfolded cylinder, we adopted horizontal periodic boundary conditions for the world. Once the cells reach the topmost cell line, they exit the simulation and are no longer considered in the model dynamics.

### 2.4. Agents

Agents represent cells divided into two physiological typologies: *transit-amplifying* cells and *differentiated* cells; and into three neoplastic typologies: *pre-adenoma, adenoma*, and *tumoral* cells. See the list of the relevant dynamic variables in Table 1 1.

**Table 1:** Cell variables

| Main Cell Variables | Values |
| --- | --- |
| Lifetime | Normal = min 96 h (hours),<br>max 120 h, Tumoral = min 96 h, max 150 h |
| Mitosis time | Normal distribution<br>mean = Mitosis mean time, variance 1 |
| Mitosis mean time | Normal (transit-amplifying) = 24 or 16 h<br>with K-ras mutations = 12 |
| Beta-cat | Descendent gradient set at 1 in the bottom<br>0 in the top of the crypt |
| Pre-adenoma | Boolean variable<br>true if both APC = [1 1] |
| Adenoma | Boolean variable<br>true if APC = [1 1] and K-ras = [1 0] or [0 1] |
| Tumoral | Boolean variable<br>true if APC = [1 1], K-ras = [1 0] and P53 = [1 1] |

Every cell, that perform autonomously the activities schematized in Fig.2, has a specific position (patch) in the world and it moves and enters into a mitosis in relationship with neighboring cells. This modeling choice is due to the possibility of preserving individual behavioral heterogeneity along with population features. The movement of cells in the physiological state has only one degree of freedom, that is, cells are allowed to move only one patch forward and only if the patch in front of them is empty, see Fig. (3, a). This procedure guarantees a constant flow of cells from the bottom to the top of the crypt. Given the fact that cells are not created with the same age, some of them die during the migration leaving empty spots in the cell line, this inconvenience is fixed enabling all cell typologies to carry out mitosis as follows: one of the cells near the empty space randomly enters mitotic activity filling the space. Another kind of inconvenience is that when a cell reaches its mitosis time it is possible that the spots near it are not empty; in this situation the mitotic cells kill the nearby cell with a lower number of mitotic activity or a higher life-time. These two processes seem to us as perfectly biologically reasonable because if a cell was killed for the number of times it has proliferated, thus related to a better or a worse fitness, or if it was killed because of its age, it is because a more efficient, or younger, cell can take its place[34].

**Figure 2.**
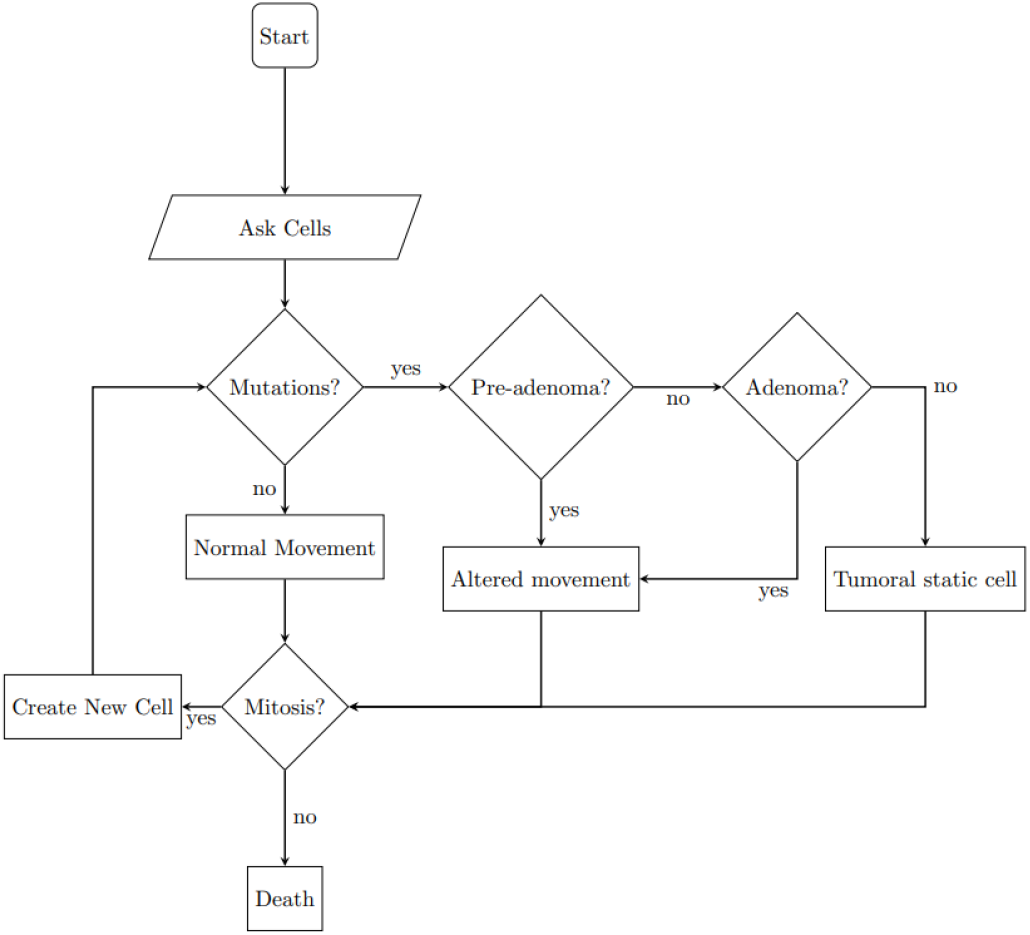
Cell behavior algorithm.

**Figure 3.**
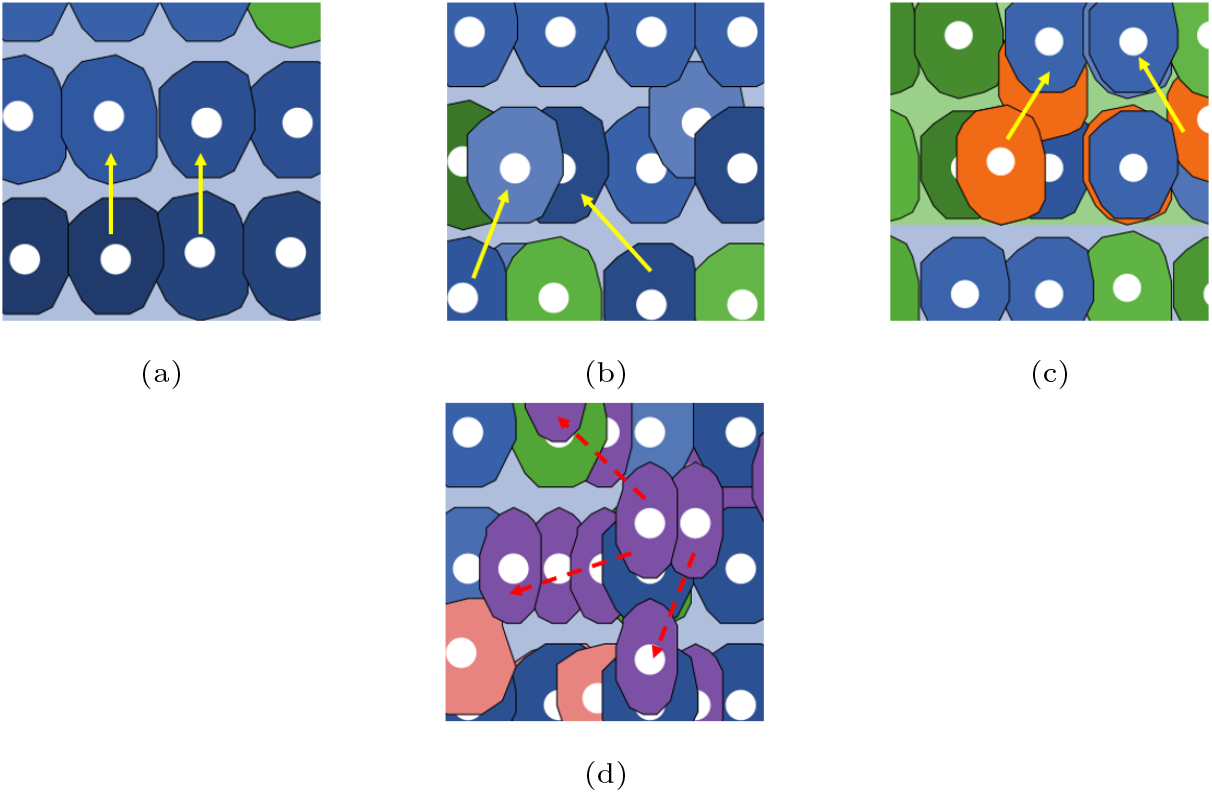
Cell movements.

### 2.5. Genetic features and mutation effects

Every cell has a set of genes, see Table 2, chosen following the CRC progression already described (see Introduction).

**Table 2:** Gene characteristics

| Genes | State |
| --- | --- |
| APC | wild type = [0 0], heterozygote = [1 0] [0 1],<br>mutated = [1 1] (positive effect in the model) |
| K-ras | wild type = [0 0], heterozygote = [1 0] [0 1],<br>(positive effect in the model) |
| P53 | wild type = [0 0], heterozygote = [1 0] [0 1],<br>mutated = [1 1] (positive effect in the model) |
| Regulation genes<br>(N = 50) | N = [ [0 0], [0 1], ..., [1 0] ]<br>determine different phenotypes of tumoral<br>cells, apoptosis, resistance to the immune system |

Each of the three genes controls a function that, in order to avoid excessive complexity, is represented as a simplified action. We assumed the one-hit hypothesis for oncogenes (*Kras* in this model) and the two-hit hypothesis for tumor suppressor genes (*APC* and *P53*) in this model), which affirm that one allele mutation is sufficient for enhancing the function of oncogenes, while a tumor suppressor gene needs both alleles to be mutated [4]. To represent another real characteristic, namely *Loss of Heterozigosity* (LOH) [35], we set the probability of point mutation in the second allele to be one order of magnitude less than the probability of point mutation in the first allele.

As a tumor suppressor gene, *APC* has a positive function when wild-type even if heterozygote, that is, it controls the differentiation of cells from *transit-amplifying* to *differentiated*. Mutation of both alleles in *APC* causes a halt in the lowering of *β*-catenin, which prevents cells from differentiating, potentially causing *transit-amplifying* cells to migrate to parts of the crypt where there should only be *differentiated*, non-proliferative, and functional cells; cells are also programmed to move not perfectly forward to simulate a slower rate of migration, which can favor cell accumulation after mitosis. The mutation of one allele in Kras increases the mean proliferation time, see Table1 ^1^.

The wild type version of *TP53* is known as the *genome Gatekeeper* [2] and while its wild type homozygote guarantees that cells with serious DNA errors (i.e., if more than half of the 50 regulation genes are mutated) die by apoptosis. The mutation of both alleles in *TP53* increases the probability of mutation in all other genes, i.e., the probability exponent decreases by 0.5 unit for each mitosis until cellular death.^2^.

Neoplastic typologies arise combining the occurrence of mutated genes non-specific temporal order, excepted for *APC* that alone stands for *pre-adenoma* typology. When a cell adopts the pre-adenoma typology, instead of moving perfectly straight, it inclines to the right or left to a random degree, see Fig. (3, b). If, in addition to *APC* mutation, *Kras* gene mutation occurs, the cell assumes *adenoma* typology, with the same movement as *pre-adenoma* cells but also proliferating at a faster rate, as shown in Fig. (3, c). The final phenotypic change is the addition of the *TP53* mutation, which causes tumoral typology by not allowing cells to move around but allowing them to proliferate all around the position rather than just adjacent spaces, see Fig. (3, d).

### 2.6. Micro-environmental features

Genetic triggers of cancer transformation are well known for many types of cancer, however, the hardest aspect to represent is the one regarding environmental factors.

Actual knowledge about the *tumor micro-environment* (TME) shows the richness of elements present in our organism from the early stages to metastasis [36], and how all these elements concur to tumor progression or tumor reduction. In order to avoid simulating every specific molecular interaction that is inconsistent with how the model is intended, we decided to design immune system cells as a combination of *natural killer cells* (NK) and *macrophages* (M1 and M2) in order to deal with TME complexity. The other environmental feature is “the population”, which simulates the fact that over-proliferating cells tend to create hypoxic niches where cells that are situated at a certain distance from the nearest blood vessel die [37]. The interaction between epithelial cells and environmental features follows these rules: Every cell in a given spot is counted, and when a threshold is reached, immune system cells appear there and move randomly. If they come across a non-physiological cell, they move toward it and kill it, making the interaction appear to be merely spatial. The same counter procedure also activates another threshold, which is the maximum number of non-hypoxic cells in a given spot.

## 3. Results

Starting values of cellular genotypes can be set to either homozygous wild-type, homozygous mutated, or heterozygotes (i.e., [0 0] and [1 1] or [0 1], [1 1]). The probability to have one of these starting genotypes can be chosen with a slider that ranges from 0 to 1 in the model interface: 0 means that all cells start with a homozygote wild-type genotype, and 1 means that all cells start with a heterozygote genotype. This characteristic is functional to the fact that we do not consider stem cells in the model dynamics, but we want to save the stem-niche hypothesis that states that mutations can also occur within stem cells, making it possible for stem clones to invade the entire crypt [28].

We chose to make several runs with three stop conditions: *a*) one year is passed; *b*) the proportion of *transit-amplifying* and *differentiated* cells inverted; *c*) crypt population is equal to 10.000 individuals. In order to observe changes in the dynamics of cell population growth, the rate of death by physiological and environmental causes (i.e., the number of cells killed by the immune system or by over-grouping causes), and the rate of death by the ageing of tumoral cells, we changed the probability of gene mutation from 10^−9^ to 10^−2^, while maintaining the other parameters constant. The other fixed parameters were:

- **number of immune system’s cells**: 1 / hour
- **number of cells that a killer cells eliminates before die**: 5 cells
- **hypoxic threshold**: 5 cells / patch
- **probability of starting genotype**: *P* = 0.5

Results have shown that all the simulations with a gene probability of mutation lower than 10^−6^ Fig.(4, A), did not present any kind of neoplastic phenomenon, and no trigger mutation occurred. Instead, all the simulations with a probability equal to 10^−6^, or higher Fig.(4, B-F), showed an exponential growth of neoplastic phenotypes, in particular adenoma and plain tumoral phenotypes, after a series of minor explosion of proliferation. The time at which the neoplastic phenotypes appear and the size of these various populations vary beteween distributions higher than 10^−6^,in fact, from 10^−5^,Fig.(4, C), to 10^−2^, Fig.(4, F)), not only are fully tumoral cells present but also *pre-adenoma* and *adenoma* typologies, with an expansion in both *transit-amplifying* and *differentiated* cell populations, see Fig. (4, F).

**Figure 4.**
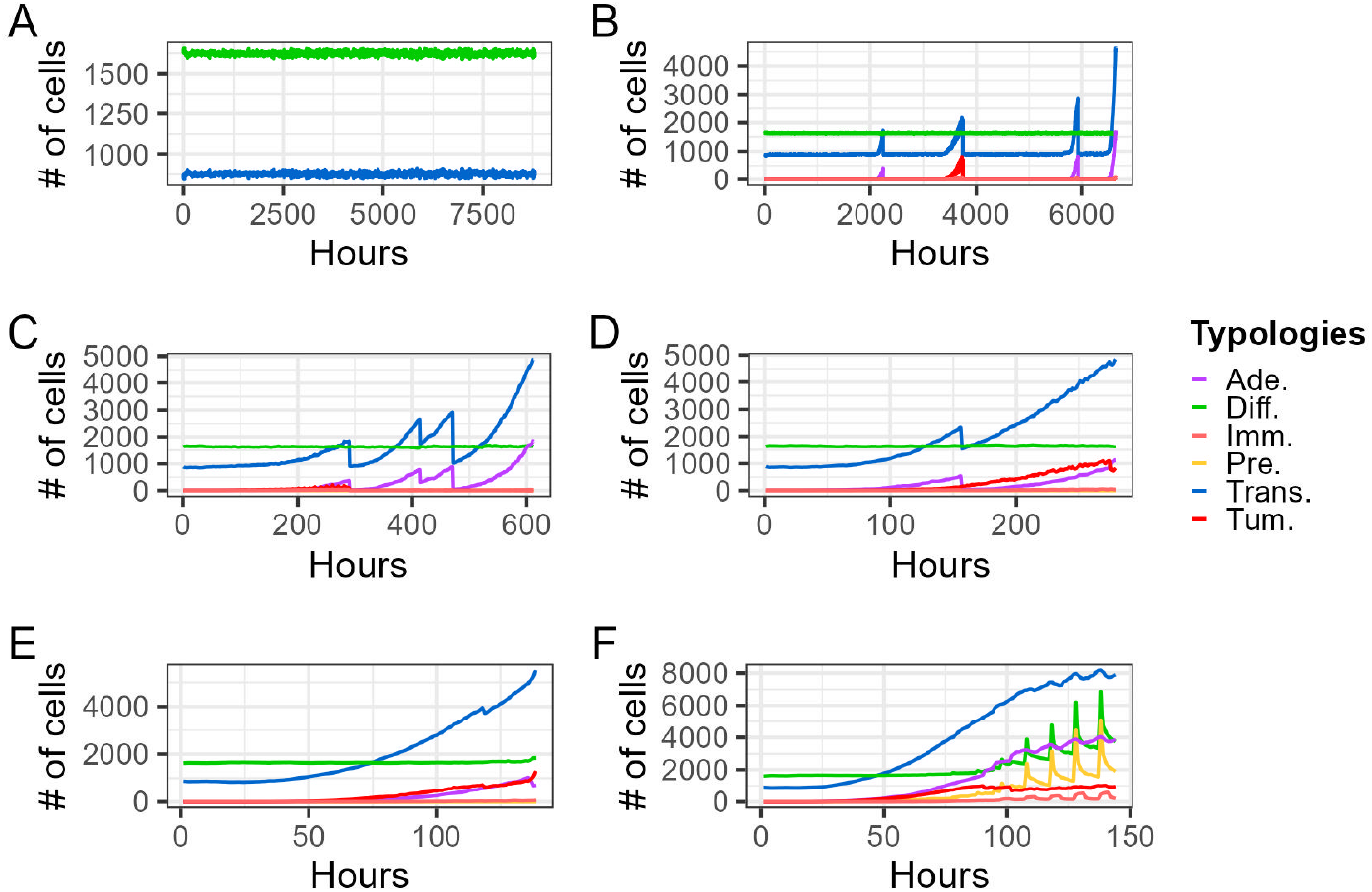
Population dynamics

Differences in death rates per hour and modes were not significant, as shown in Figure (5, A), but they became evident at a death rate of 10^−4^. In fact, we observed a decline in the rate of normal deaths and a corresponding rise in other death modalities, with the rate of hypoxic deaths being slightly higher than the death rate caused by immune system cells, (5, B).

**Figure 5.**
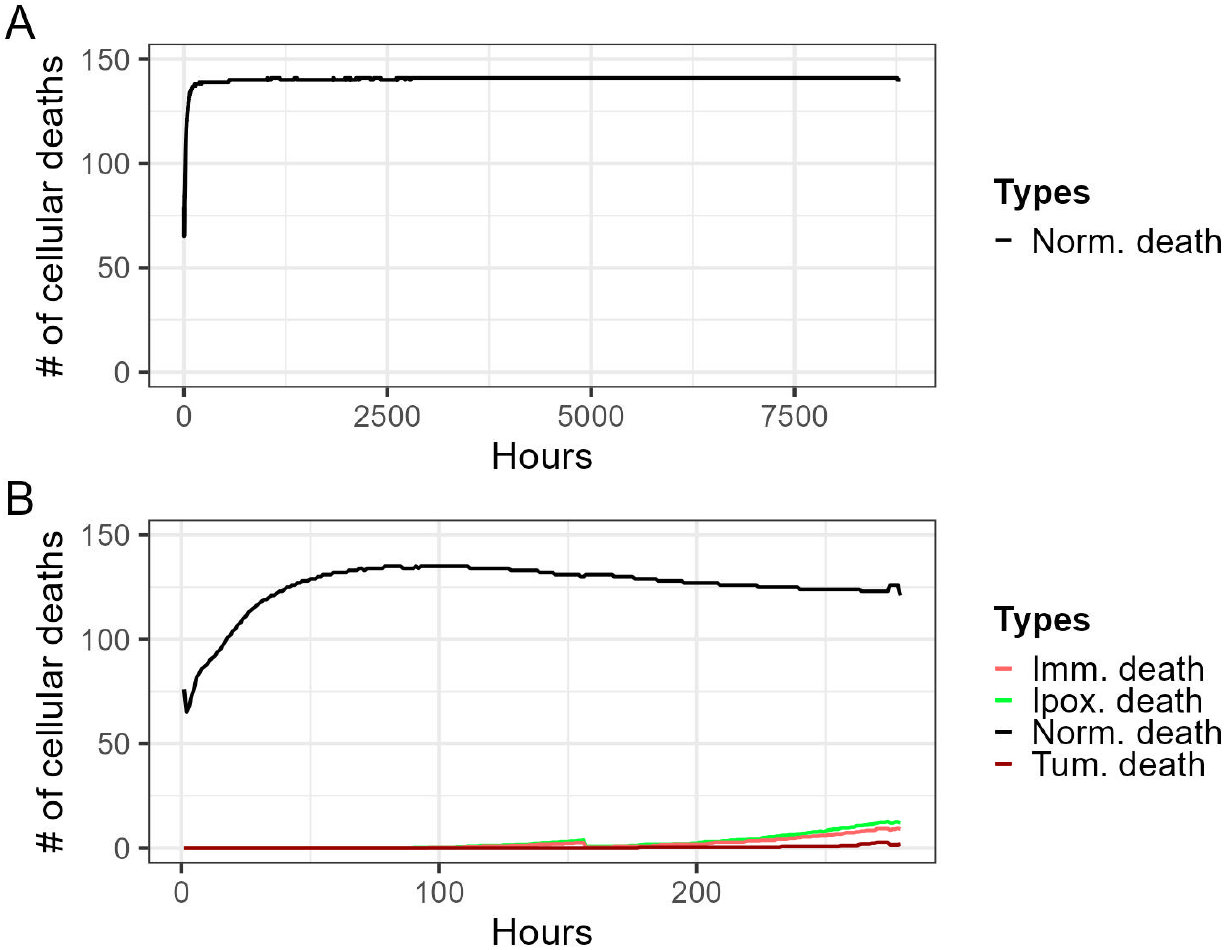
Cells death types

The results of genotype frequencies revealed a satisfying visual correspondence between the flow of genes within populations and the dynamics of the population. In fact, Fig. (6, A) depicts a constant genetic flow, mirroring the constant dynamics of the population at 10^−9^, while Figures (6, B) and (6, C) show the corresponding dynamics of the population at 10^−6^ and 10^−2^. Interesting results emerged concerning the number of proliferation events for each phenotype. In fact, normal cells seem to have a skewed normal distribution, Fig. (7, A), while *adenoma* and *tumoral* cells showed a very wide normal distribution with a nearly shared mean value, see Fig. (7, B-C).

**Figure 6.**
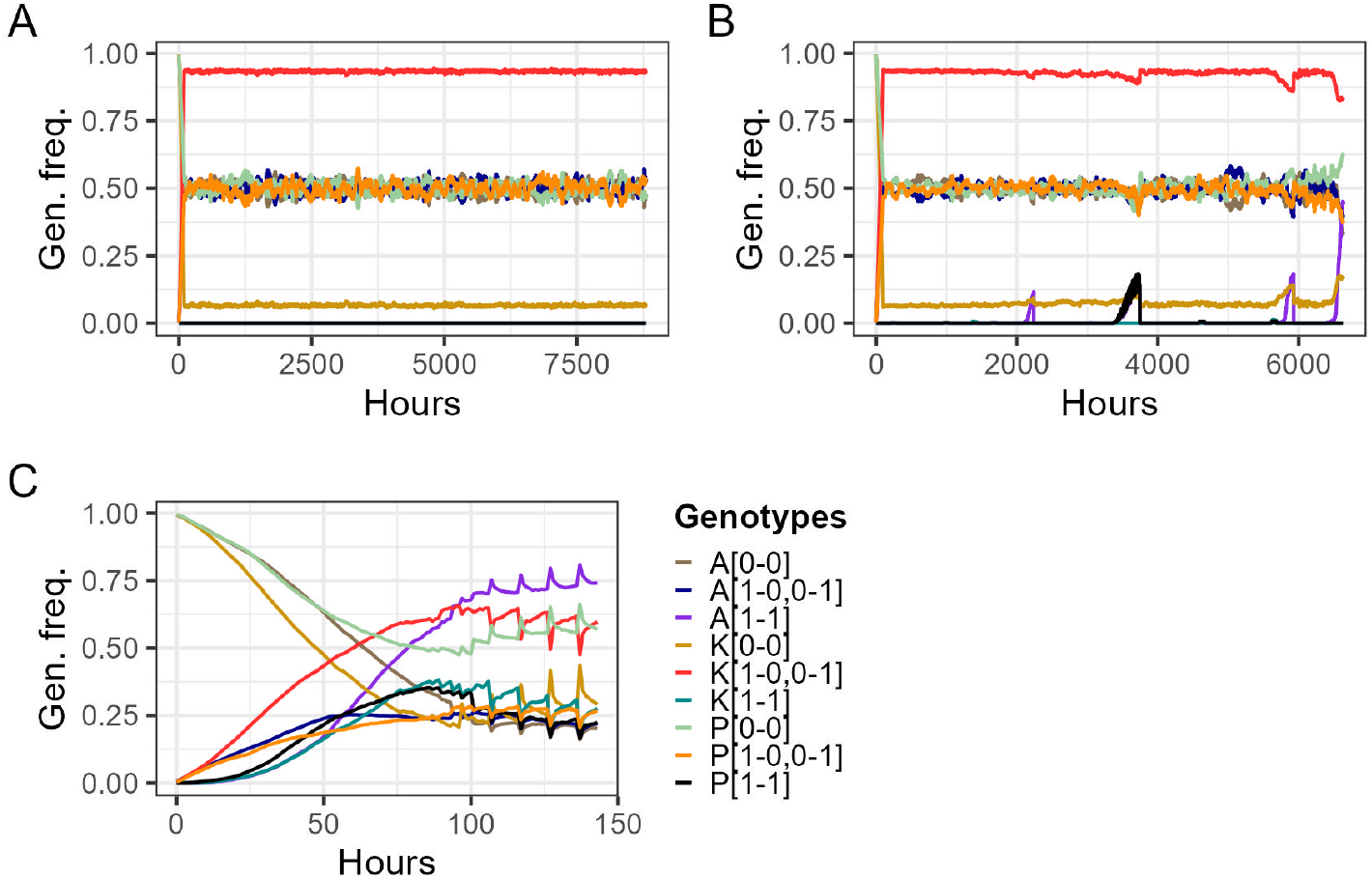
Genotype frequencies

**Figure 7.**
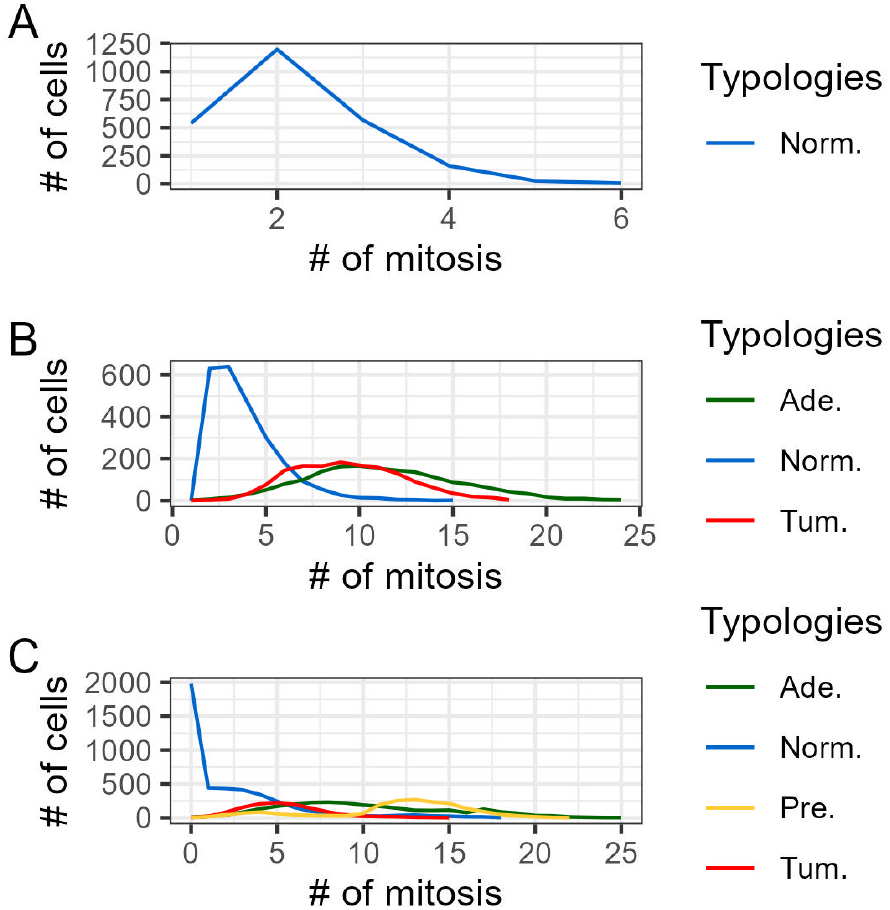
Proliferation distributions

The other relevant feature was the density of cells that increased in value up to 7 cells per spot, for a probability of mutation higher than 10^−7^, see Fig. (8).

**Figure 8.**
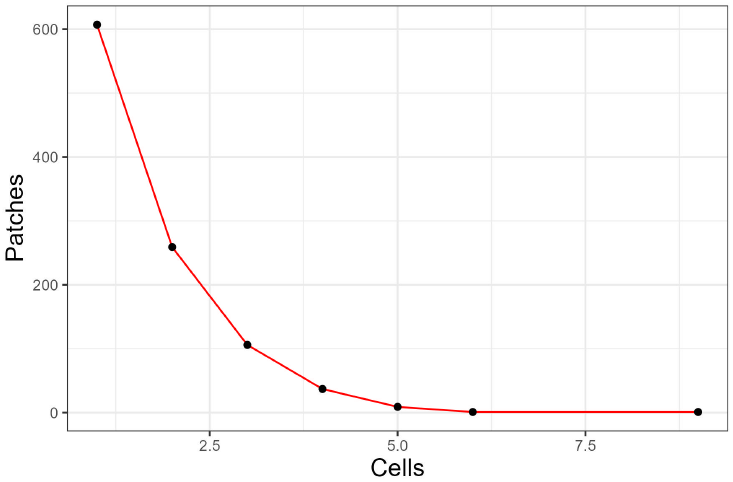
Cells per patch

### 3.1. Discussion

In building up this model we meant to create a device to show theoretical investigations, make hypotheses, and design experiments about a biological phenomenon. However, a model constructed with the aid of data and assumptions drawn from the literature should provide results that are, at least, not in contradiction with the state of the art. In our opinion this is clear at least from three points of view: the first one is visual, by the fact that the world of Net-logo shows how cells are arranged in the crypt giving rise to the phenomenon called *Intra-tumoral Heterogeneity* (ITH) [38], see Fig. (10, D-F); the second one is quantitative, based on how the model is conceived, features such as proliferation events measured by crypt height, are similar to those of other CRC studies [39], see Fig. (9); the third one is that the model follows the tumor development as described by Fearon and Vogelstein in their famous work [3]. Visual aspect shows the enormous difference between crypts at 10^−9^, see Fig. (10, a) and crypts at 10^−6^, or higher probability of mutation, see Fig. (10, B-F). When we compare population dynamics in the range (10^−9^-10^−6^), what we observe is something like a phase transition, moreover, if we look at the range (10^−3^-10^−2^) we observe the behavior attributed to cells with CIN (*chromosomal instability*) [35], with a rapid growth of total cellular population (4.000-6.000 individuals) in 100-150 hours. Some remarks are necessary about the initial genotype condition of the cell population, in fact, all the runs were carried out with the starting probability of heterozygosis set up on (*P* = 0.5), and it is not reported that, for every generation cycle, 50% of stem cells could have a mutated allele. Therefor, it is necessary to conduct tests with other starting values. Immune system cells and over-grouping deaths are the other parameters that need further tests, at least the one that consists in a minimal genetic model of tumor development, that is, the number of immune cells set to 0 and the hypoxic threshold set to a large number, in this scenario we should observe a distinct tumor development once the first neoplastic cell arises. Coherent with the Vogelstein model, we observed that in all the simulation where neoplastic phenotypes arose, during the early stages of the tumor growth, cells first assumed the *pre-adenoma* phenotype (i.e., *APC* mutation) or at most the *adenoma phenotype* (probably because *Kras* act as an oncogene even with one mutated allele, and genotype frequencies show that almost the entire population is heterozygote for *Kras*). From here, we can develop a potential scenario. Imagine a *pre-adenoma* cell that grows and spreads throughout the crypt, interfering with the growth and movement of other cells, causing them to die or become immobile. Then, imagine that the *pre-adenoma* cell acquiring another mutation, becomes an *adenoma* cell that grows at a rate twice as fast as the others, creating a mass of cells. Finally, imagine that this mass of cells reaches a fatal threshold, which kills cells of every type in a given radius, but the ones that can proliferate more than the others have a selective advantage and continue to proliferate and grow in the crypt. These cells also have the ability to avoid the action of immune system cells on a spatial level, which causes them to develop additional mutations that turn them into tumors. In this final stage, they remain static, proliferate at an even faster rate, and die later than the physiological cells, bringing the entire crypt to a fully *tumoral* and metastatic state.

**Figure 9.**
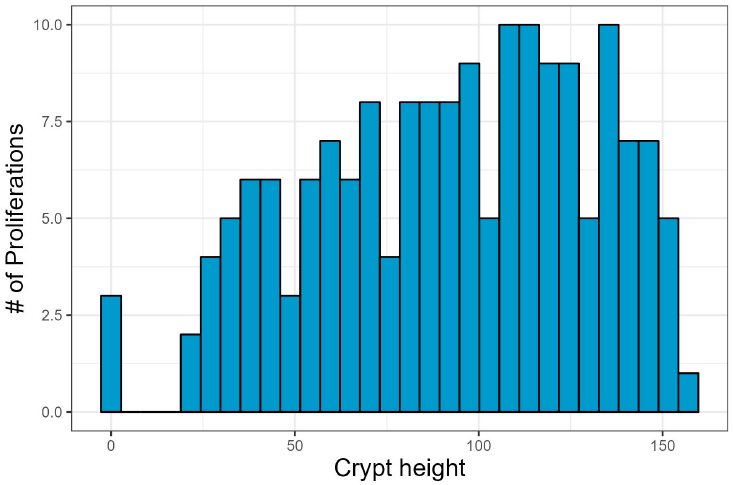
Mitotic activity based on crypt height.

**Figure 10.**
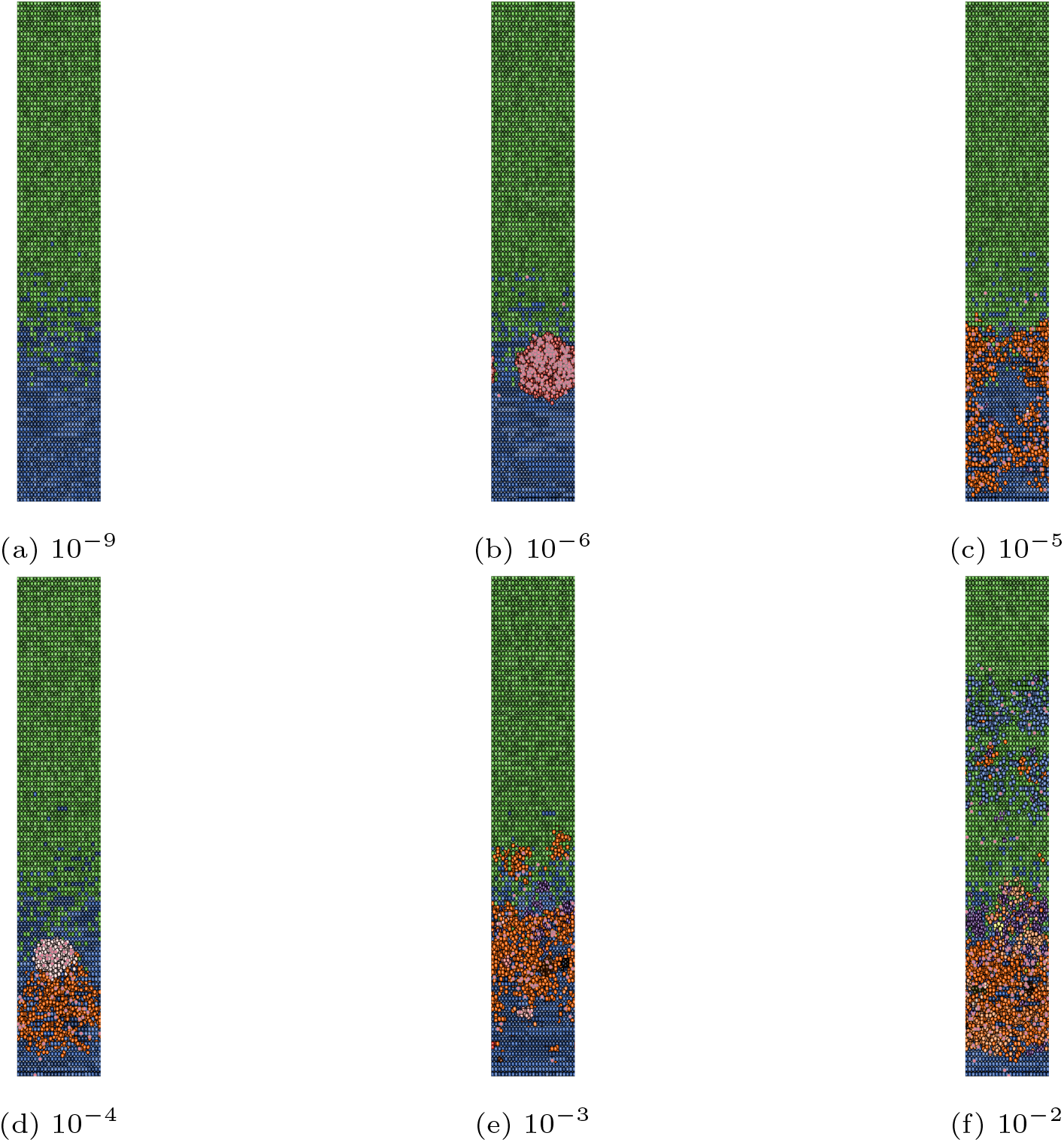
Crypt final states

## 4. Conclusions

In this work, we try to introduce a new type of model that takes into account both genetic and environmental components. **SMT** (*somatic mutation theory*) and **TOFT** (*tissue organization field theory*) are the two main paradigms used in cancer research that represent these two aspects. The first one assumes that multiple consecutive gene mutations in one cell, which proliferate more than the others, spread the tumoral phenotype and are the primary cause of tumor emergence and development. The second states that environmental factors are to blame for these alterations in the signaling between cellular populations in adjacent tissues, which results in gene mutations [40, 41].

The connection between these two aspects, even if it is oversimplified, is built in a way that is consistent with what cancer research indicates, namely that it is not just a matter of mutation and hyperproliferation. The presence or absence of immune system cells favors the rise of one cell typology over another, and an elevated number of cells in a small space causes cells to die. Additional research should provide more precise evidence of correlations (and hopefully causal relationships) between system variables and environmental factors that affect the dynamics of different cell typologies.

## Footnotes

1 We assumed that the distribution of mitotic time is well represented by a normal distribution, which is typical, for some features, in biological populations.

2 The decrease amount is not taken from the literature, to our knowledge there is no quantitative analysis about such a phenomenon, and is chosen to be not particularly prominent nor insufficient simulating DNA deregulations.

